# Quantification of discrete gut bacterial strains following fecal transplantation for recurrent *Clostridioides difficile* infection demonstrates long-term stable engraftment in non-relapsing recipients

**DOI:** 10.1101/2020.09.10.292136

**Authors:** Varun Aggarwala, Ilaria Mogno, Zhihua Li, Chao Yang, Graham J. Britton, Alice Chen-Liaw, Josephine Mitcham, Gerold Bongers, Dirk Gevers, Jose C. Clemente, Jean-Frederic Colombel, Ari Grinspan, Jeremiah Faith

## Abstract

Fecal Microbiota Transplantation (FMT), while successful for the treatment of recurrent *Clostridioides difficile* (rCDI) infection, lacks a quantitative identification of the discrete bacterial strains that transmit and stably engraft in recipients, and their association with clinical outcomes. Using >1,000 unique bacterial strains isolated and sequenced from a combination of 22 FMT donors and recipients, we develop a statistical approach *Strainer* to detect and track sequenced bacterial strains from low depth metagenomic sequencing data. On application to 14 FMT interventions, we detect stable and high engraftment of ∼71% of gut microbiota strains in recipients at even 5-years post-transplant, a remarkably durable therapeutic from a single administration. We found differential transmission and engraftment efficacy across bacterial taxonomic groups over short and long-time scales. Although ∼80% of the original pre-FMT recipient strains were eliminated by the FMT, those strains that remain persist even 5 years later, along with newer strains acquired from the environment. The precise quantification of donor bacterial strains in recipients independently explained the clinical outcomes of early and late relapse. Our framework identifies the consistently engrafting discrete bacterial strains for use in Live Biotherapeutic Products (LBP) as a safer, scalable alternative to FMT and enables systematic evaluation of different FMT and LBP study designs.

## Introduction

Several studies have demonstrated that gut microbiota strain variation impacts health^1–7^ through mechanisms including altered immune function^8,9^ and drug metabolism^10,11^. Manipulation of gut microbiota strain composition therefore provides a potential route to influence health^7,12–15^. Clinically, the dominant therapeutic application of gut microbiota additive engineering is Fecal Microbiota Transplantation^16^ (FMT) for recurrent *Clostridioides difficile* infection (rCDI), which has success rates of over 90% across a variety of administration routes^17–21^ with documented but much lower efficacy for ulcerative colitis^22–24^. However, our understanding of FMT dosage, study design, predictors of response, and mechanism of action is still in its infancy^25–27^. Since the functional impact of the gut microbiota is at the level of strains, quantification of gut microbiota at this resolution is essential for understanding the therapeutic potential of FMT and its impact on the health and disease of the host.

Achieving strain level resolution has been challenging because human gut microbiota comprises of numerous bacterial strains in each species^28^, the majority of which have not been isolated or even metagenomically detected before. As a result, previous microbiome analyses^29^ have largely focused on species or lower level of resolution because finding delineating features of discrete bacterial strains, a necessary step for FMT strain tracking, remains unresolved. Recent FMT analyses have combined deep metagenomic sequencing with the computational detection of single nucleotide polymorphic (SNP) variants in common species marker genes to quantify some properties of strain composition. These approaches^30–33^ have found enrichments of donor SNPs in the recipient microbiota post-FMT suggesting transmission and engraftment^31,32^ of some proportion of the donor’s strains in the recipient but linkage of these donor microbe SNPs to the donor’s discrete bacterial strains remains elusive. While informative, these approaches require very deep metagenomic sequencing to track strains present at even shallow relative abundance^34^. In addition, they do not accurately model the microbiota as a defined and finite set of strains. This linking of bacterial genetic variation into a discrete unit, i.e., a cultured bacterial isolate, is essential for understanding transmission and for fulfillment of the commensal Koch’s postulates^35,36^.

Recent advances in high throughput bacterial culturing have enabled the isolation of significant fraction of the gut microbiota of an individual^37–42^. Sequencing the genomes of these isolates enables the tracking of bacterial strains by whole genome comparisons^1,39^ with extremely high resolution but low throughput. A hybrid approach between metagenomics and culturing-based strain tracking is to first comprehensively culture and sequence the strains from a limited number of samples in each individual of interest and then track each strain across one or more metagenomic samples in which it might appear using the strain’s genome sequence. Such an approach can track strains as a single linked entity and is more sensitive than inferring SNPs in marker genes, as confidence in the detection of a strain within a metagenome is determined by the combined presence of multiple unique SNPs or k-mers across a genome.

Here we use metagenomic sequencing, combined with bacterial strain culturing from the fecal microbiome of FMT donors and rCDI recipients, to precisely quantify the engraftment and stability of strain transmission from donors to recipients. To do so, we develop and experimentally validate a statistical algorithm *Strainer*, that detects bacterial strain genomes at high precision and recall from metagenomic sequencing data. We apply *Strainer* to track 1,008 unique bacterial strains isolated from donors and recipients across 85 metagenomics samples from rCDI recipients to estimate strain transmission and long-term engraftment. We find the majority of donor strains engraft in non-relapsing rCDI FMT recipients. Low donor engraftment explained relapse and donor engraftment explained success in patients at all timepoints. Importantly, we find the majority of engrafted strains from the donor are long-term stable for the entire 5-year sampling period in the recipient gut. These results suggest FMT represents a semi-permanent alteration of the host microbiome – a remarkably durable therapeutic from a single administration, whose stability resembles that of healthy controls^39^.

## Results

### Developing and experimentally validating a strain detection algorithm, *Strainer Strainer* algorithm

The central challenge behind strain tracking from metagenomics data, is the identification of a set of informative sequence features or k-mers from the bacterial genome that can be detected in multiple individuals. Since the field is far from sequencing the majority of bacterial strains and each species contains numerous closely related unique strains which share a majority of genomic content^39,43,44^ (in case of *Bacteriodes ovatus* 54%, sd = 16%, **Supplementary Figure 1A**), identification of such informative features to track strains is a challenge. To obtain the most informative k-mers for tracking a given strain (overview of our algorithm *Strainer* in **Supplementary Figure 1B**), we first removed k-mers (k = 31) shared extensively with other strains from a set of >100K bacterial genomes from NCBI, >1000 bacterial genomes from this study and in any sequencing read from an independent set of 110 unrelated metagenomes^45^. The number of k-mers eliminated varied highly across species (**Supplementary Table 1, Supplementary Figure 1C**), suggesting the strains in some species share more genomic content. We also processed each sequencing read in a metagenomic sample, and if it had k-mers belonging to multiple strains or a high proportion of non-unique k-mers identified, suggesting all k-mers in the read are likely non-informative, we removed all of them from the initial set for this strain. We next assigned to each strain the metagenomic sequencing reads, from a given sample of interest, that have a dominant proportion of informative k-mers (more than 95% of total). We then mapped these strain-assigned reads to the corresponding strains’ genome and quantified only the non-overlapping reads. This step adjusts for evenness of coverage at the genome with the assumption that metagenomic reads should be randomly distributed across the genome for true positive strains colonized in the host’s gut. Finally, we compared the non-overlapping reads for a strain in the metagenomic sample, with those found in negative controls (non-cohabitating and distant samples where probability of occurrence of the same strain is very low^4,39^) to find the enrichment of informative reads and assign a confidence score for presence of the strain in the sample.

### *Strainer* validation on a defined community of strains in gnotobiotic mice

We next tested the ability of *Strainer* to detect bacterial strains in situations of varied strain complexity, by colonizing gnotobiotic mice with a subset of 10 unique strains of the common human gut commensal bacterium *Bacteroides ovatus*. These mice were either monocolonized or colonized in the context of defined culture collections of bacteria isolated from 3 different human fecal samples^7^ (**Figure 1A, Supplementary Table 2**). Our goal was to accurately detect the set of *B. ovatus* strains in each mouse from its fecal metagenome. We successfully detected strain F and only strain F in all fifteen mice in which it was colonized in the context of human cultured microbiome library 1 which contained strain F (**Figure 1A**). When we increased the *B. ovatus* strain complexity of the microbiome by colonizing the gnotobiotic mice with both a human culture library and a set of 4 or 8 different strains, we found that strains administered to the mice were only detected from the corresponding metagenome. We quantified our overall performance in these simplified communities using precision and recall, which were 100% and 86.9% respectively, with no false positives in 280 different tests (specificity 100%).

**Figure 1.**
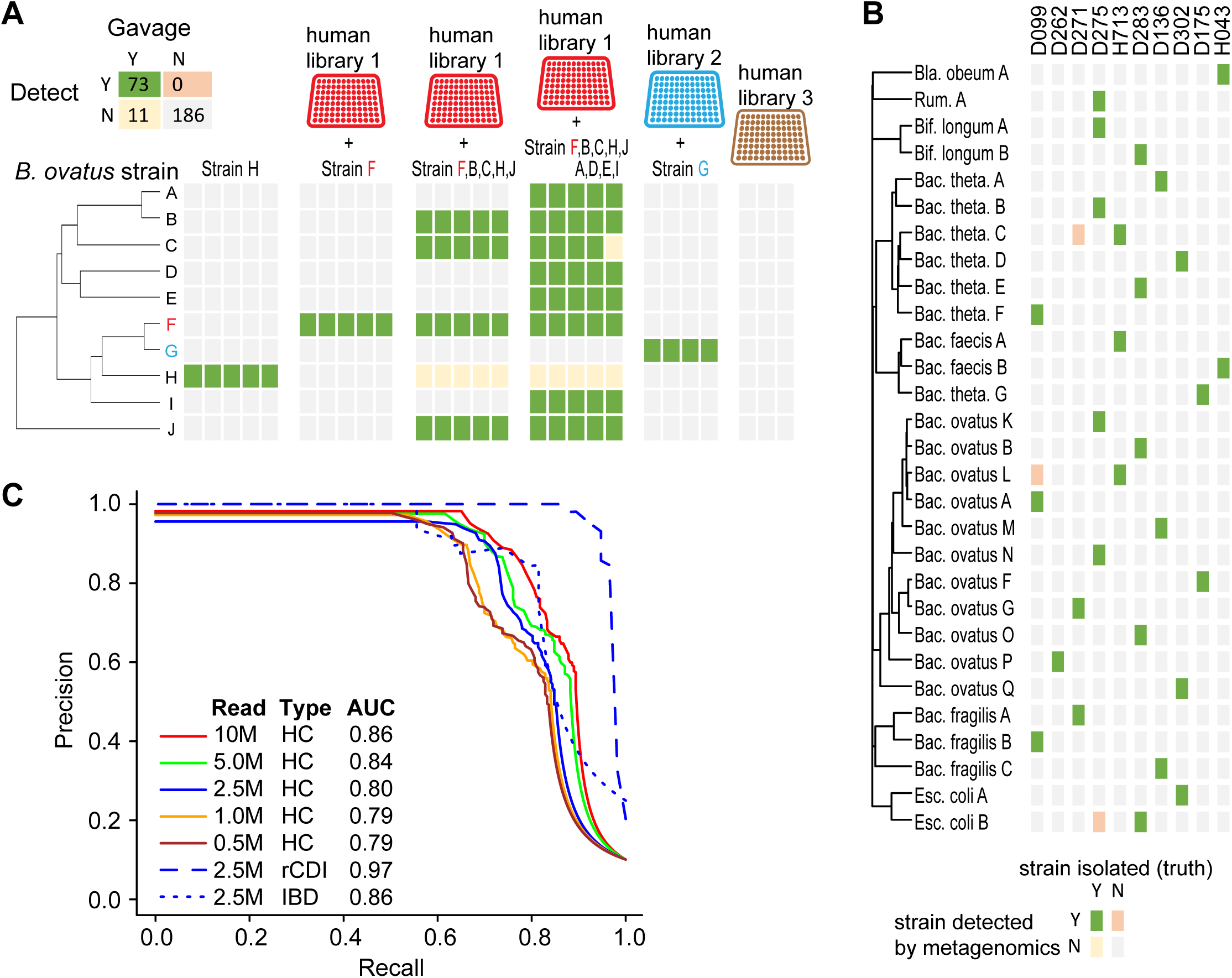
*Strainer* algorithm accurate detects bacterial strains from complex gut communities through low depth metagenomics. (**A**) *Strainer* can accurately detect the correct *Bacteriodes ovatus* strain(s) in gnotobiotic mice, from other closely related strains. Each column represents an independent germ-free mouse gavaged with the specific B. ovatus strain(s) with or without a diverse human gut bacterial culture library of strains. Strains F and G were contained in human culture library 1 and 2 respectively. Human culture library 3 contained no *B. ovatus*, while the remaining *B. ovatus* isolates were isolated from other human fecal samples. Green box indicates the strain was introduced in the mice and detected in metagenomics (true positive), Grey indicates the strain was not detected and (true negative), Orange indicates the strain was detected but was not introduced (false positive) and Yellow indicates the strain was not detected but was gavaged in the mice (unknown as gavaging a strain does not always lead to stable colonization). (**B**) Representative examples highlighting our algorithm’s ability to match strains to the correct human fecal sample for closely related and evolutionary distant taxa. Green box indicates the strain was isolated from the human fecal sample and detected by our algorithm (true positive), Orange indicates the strain was detected but not isolated from the sample (false positive) and Grey indicates that the strain was not detected and not isolated from this sample (true negative). (**C**) Performance assessment of *Strainer*’s ability to match strains to the metagenome of the sample from which they were isolated. Solid lines denote the results at different sequencing depth after application of our algorithm on 261 strains isolated from healthy controls (HC). The color blue indicates the sequencing depth of 2.5M reads, while the dashed line indicates the result after application of *Strainer* on 56 strains isolated from patients with rCDI and the dotted curve is for 54 strains from patients with IBD. AUC of the Precision-Recall curves is in the legend box.

### *Strainer* validation on complex human gut microbiotas

We next tested our strain detection approach in the context of several complex human gut microbiota communities with high species overlap but little to no strain overlap. This resembles the use-case application for FMT where a potentially transmitted bacterial strain has to be precisely detected across multiple individuals, while differentiating it from other related commensal strains from the same species. We sequenced the fecal metagenome of 10 unrelated individuals as well as the genome of 261 bacterial strains isolated from the same fecal samples (**Supplementary Table 3**). We then evaluated the ability of *Strainer* to detect each of 261 strains in the correct individual’s metagenome, while not falsely detecting it in the other nine metagenomes. Our approach worked well across varied strain complexity (i.e., one or more strains per species) and correctly matched strains within and across species to the correct sample from which they were isolated (**Figure 1B**). With 10M metagenomic reads per sample, we reached a precision of 100% at a recall of over 60% with an AUC of 0.86 (**Figure 1C**). Although we attained slightly higher recall with deeper metagenomics, even 500K metagenomic sequencing reads were sufficient to reach precision of 100% at a recall of over 50% with an AUC of 0.79 (**Figure 1C**). Given that our goal in this study is to detect strains in individuals with rCDI and the increased prevalence of rCDI in subjects with Inflammatory Bowel Disease (IBD), we generated similar testing datasets (**Supplementary Tables 4-5**) from five individuals with rCDI and four individuals with Inflammatory Bowel Disease (IBD). We found similar results between the healthy, IBD, and rCDI strain detection test sets at 2.5M reads with slightly higher AUC for rCDI likely as a result of the low diversity of the gut microbiome in rCDI^46,47^ (**Figure 1C**). Altogether these results demonstrate that the Strainer algorithm can precisely and sensitively track sequenced bacterial strains in a metagenome thus providing the potential to quantify discrete donor strain transmission and recipient strain persistence in FMT over time.

### Proportional engraftment of donor strains varies by rCDI FMT outcome

#### FMT samples and isolation of strains

To apply our validated strain detection algorithm to understand short and long-term engraftment of donor strains and persistence of recipient strains in the context of FMT for rCDI, we isolated and sequenced 1,008 unique bacterial strains (207 species) from 8 FMT healthy donors and 14 rCDI FMT recipients (**Table 1** and **Supplementary Tables 1, 6-7**). Similar to our previous analyses^12,39,48^, bacterial isolates with <96% whole genome similarity were defined as unique strains, otherwise they were considered as multiple isolates for the representative strain. In parallel, we sequenced an average of ∼5.2M metagenome reads from each donor fecal sample used for the transplant and of recipient fecal samples taken prior to and for up to 5 years after FMT (85 metagenomic samples). As in prior studies^42,49^, these cultured strains represented the majority of the metagenome with 70% (sd = 16%) of bacterial metagenomic reads mapping to the cultured strain genomes (**Supplementary Figure 1D**). Contig building on the remaining bacterial reads did not generate any large contigs, and the average length after combining all contigs was only 1.6 million base pairs, some of which likely results from sequencing noise. We also evaluated the comprehensiveness of our cultured bacterial strain library by gavaging several germ-free mice with human stool and performing metagenomics on the mouse fecal samples. The cultured bacterial strains which explained 78% of bacterial reads from the human metagenomics sample, explained up to 97% of bacterial reads in the gnotobiotic mice with the same human stool, suggesting the majority of unexplained bacterial metagenomics reads in the human sample were from unculturable sources (e.g. dead bacteria from food and environmental sources).

**Table 1.**
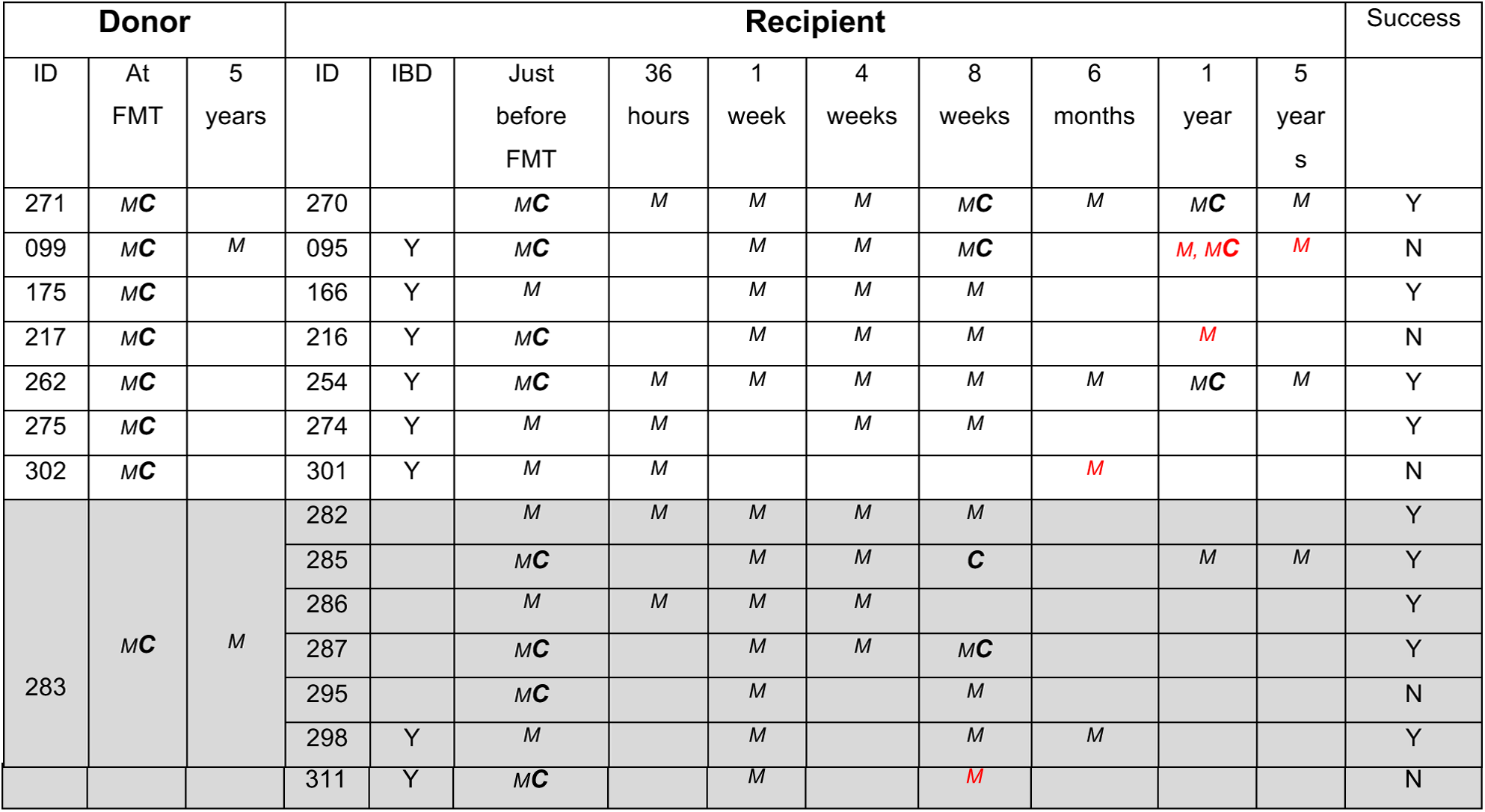
Samples, metagenomics, and culturing available from each donor and recipient. 8 donors provided their fecal material for FMT to 14 patients with either rCDI or both rCDI and IBD. Fecal metagenomics was performed on all stool samples. Donor strains from all the donors were isolated and tracked in matching recipient metagenomes over time. Strains were also isolated from a few recipients both pre- and post-FMT. *M* and ***C*** indicate that metagenomics or culturing respectively were performed at an indicated time point. The red highlight denotes that a sample was collected after repeat FMT (due to initial failure of FMT). Success indicates that no relapse was noted for that patient.

#### Early engraftment in FMT recipients

In the clinical cohort, seven FMT donors each provided their sample to a single recipient, while one donor provided the sample for seven different patients (**Table 1, Figure 2A**)). Since many FMT interventions evaluate the clinical outcome at 8-wks post-FMT, we used *Strainer* to measure the engraftment of donor strains in the recipients at this timepoint to evaluate if it can independently explain outcome. We defined the Proportional Engraftment of Donor strains (PED) as the number of donor strains detected in a recipient post-FMT divided by total number of strains isolated from the donor. Mean PED at 8 weeks was 72% across all 11 rCDI individuals with no early relapse within 2 months post-FMT (**Figure 2B**). We did not find a difference in PED between non-early relapsing rCDI recipients with IBD vs no-IBD (**Figure 2C**, p-val=0.13 from one side Wilcox test). We did however find significantly reduced PED in patients who had an early relapse within 2 months of FMT (**Figure 2B**, p-val=0.02 from one side Wilcox test) compared to patients with no-relapse after FMT. This suggests that precise engraftment of donor strains in recipients can independently explain the early clinical outcome of an FMT intervention, as subjects could be perfectly classified into relapse or non-relapse with a PED threshold of 0.35.

**Figure 2.**
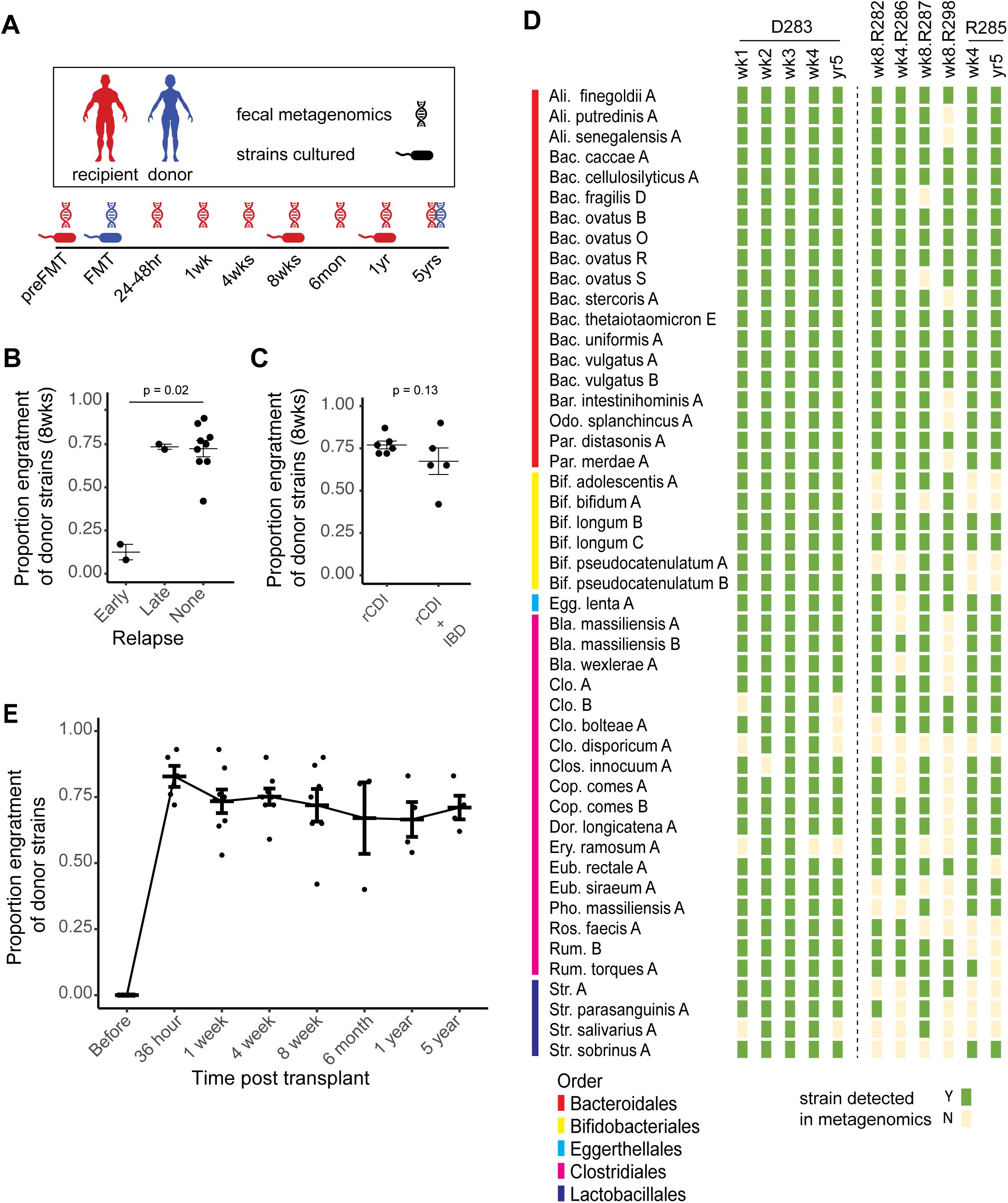
The majority of FMT donor strains durably engraft. (**A**) Overview of FMT study design indicating the dates of metagenomic sequencing and bacterial strain culturing. The genome sequences of the cultured bacterial strains are used to track each strain across metagenomic samples using *Strainer*. (**B)** Proportional Engraftment of donor’s (PED) strains at 8-weeks can predict early relapse of FMT in patients with rCDI. (**C**) Proportional engraftment of donor’s strains at 8-weeks in patients with successful FMT is not significantly different between individuals with rCDI alone and those with both rCDI plus IBD. (**D**) Bacterial strain engraftment in a single donor to multiple recipients setting. The first 4 columns are weekly metagenomic samples from the donor, while the 5^th^ column is the donor sample from 5 years later. The next 6 columns are from the FMT recipients that did not have an early relapse. The last column is from one of the recipient 5 years later. *Strainer* was used to find the presence (green) or absence (yellow) of each bacterial strain from the corresponding metagenomics sample. (**E**) Strains from the donor remain stably engrafted in successful post-FMT patients at least 5 years after transplant.

The isolation and sequencing of the transmitted strains still represents a gold standard validation, first established by Koch’s postulates for microbial pathogens^35,36^. This approach has been applied in FMT for rCDI in a limited manner using selective culturing to definitively demonstrate the transmission^50^ of three potentially procarcinogenic species. However, there is a paucity of larger studies demonstrating transmission of donor bacterial strains from multiple species and across different FMT interventions. Here, we also cultured strains from 6 recipients both pre- and post-FMT (**Figure 2A**) and compared the strain composition to that from the donor, to experimentally validate bacterial strain transmission. We never isolate a donor strain in any recipient prior to transplant, yet we isolated 48 donor strains in recipients post-FMT, encompassing 16 different species. The large majority (96%, **Table 2**) of these gold standard culture-based strain transmission determinations were also detected in the relevant metagenome (100% if we include detection at earlier timepoints) by the *Strainer* algorithm, further strengthening the confidence in our algorithmic approach.

**Table 2.**
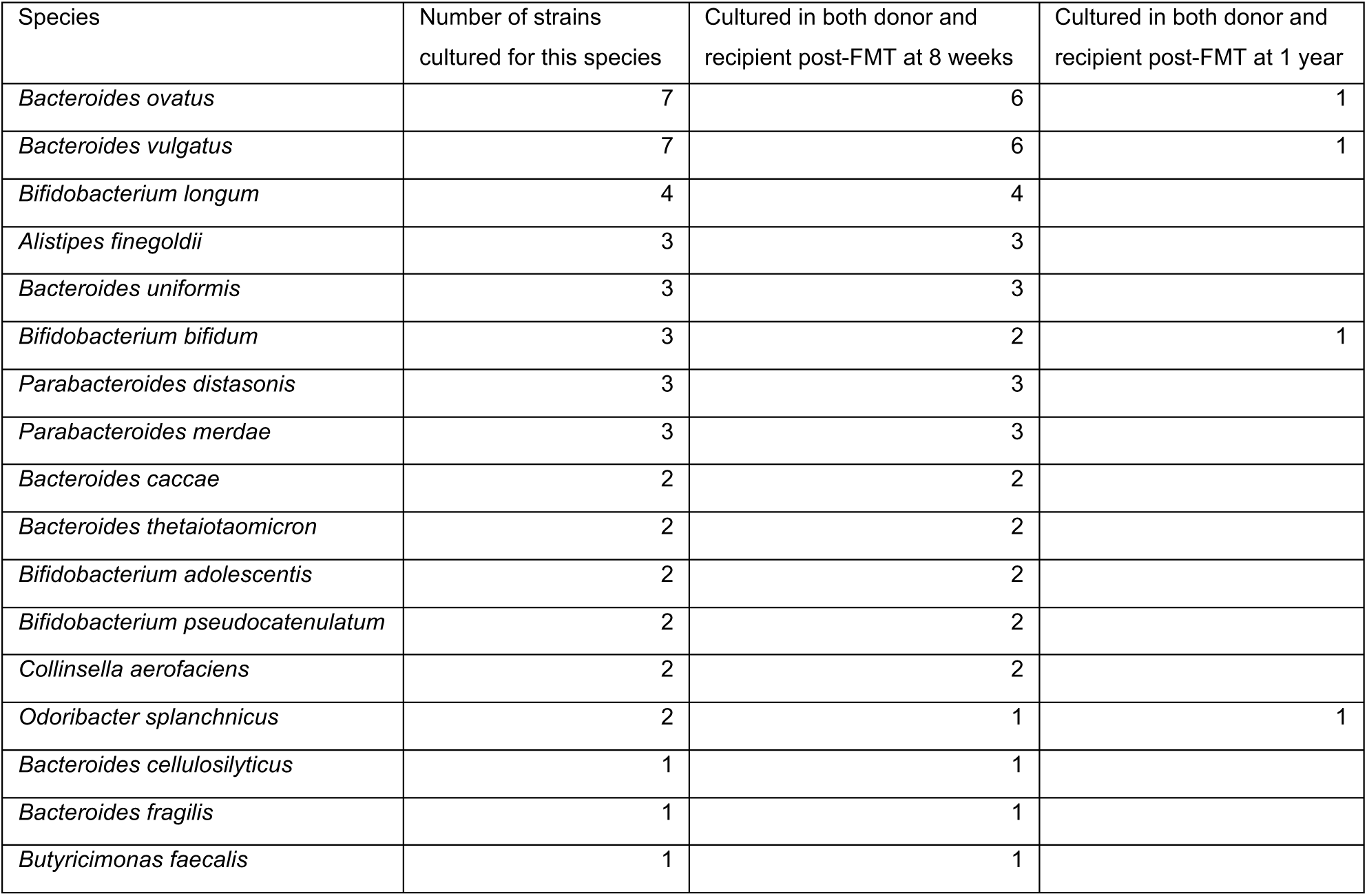
Gold standard set of bacterial strains cultured and isolated independently both from the donor and recipient post-FMT demonstrating transmission. The vast majority of these (46/48) strains were also detected independently in metagenomics samples from the same timepoint when they were cultured, and the other 2 were detected at an earlier timepoint, highlighting the *Strainer* algorithm’s capability to track and study engraftment of strains post-FMT.

### Taxa of donor strains engraft differentially in FMT recipients

As we better understand the functional impact of gut microbiota strains on health, knowing the engraftment potential of these strains might inform future FMT trials, donor selection for different indications, and the development of defined Live Biotherapeutic Products (LBP)s^51,52^ as a scalable, safer alternative to FMT^31,32,53^. Engraftment of donor strains was high in non-relapsing rCDI FMT recipients but it is unknown if all taxa engraft equally well. Towards this, we first investigated the strains that transmit and engraft for 8-weeks in 4 non-relapsing recipients which had separate donors (**Supplementary Figure 2A**). Strains belonging to order Bifidobacteriales engrafted less (58% of strains), while order Bacteriodales engrafted higher (93% of strains, **Supplementary Figure 2B**, p-val < 0.008 from fisher-exact test). Next, we investigated differential engraftment in the single donor to multiple FMT recipient setting, where we have higher power to detect stable engraftment of a strain in multiple recipients. We focused on the highly transmissible strains that stably engraft in at least 4 out of 5 non-relapsing recipients from this single donor, and again found those belonging to order Bacteriodales always engrafted (100%, 19/19, **Figure 2D**) more than those from order Bifidobacteriales (50%, 3/6). Lactobacillales were never found to stably engraft in recipients (0/4). Overall, we find high engraftment of donor microbes with some broad taxonomic groups engrafting more consistently than others across several FMT interventions and study designs.

### Long term stability of strains

Given the stable engraftment of donor microbes for up to 8-weeks post-transplant and the lack of relapse in the majority of the FMT recipients, we hypothesized the newly established gut microbiota in these individuals would also be stable at longer time-scales, similar to that demonstrated in healthy human gut microbiotas ^1,37,39,54–56^. To determine if our *Strainer* algorithm could successfully detect long-term gut microbiota stability in healthy individuals, we profiled the strain stability in a healthy FMT donor over a 5-year time interval. One donor was also profiled weekly for 4 time points (Donor D283 which provided the material for FMT to several recipients, **Figure 2D**). Over this short time scale a large majority (47/48) of strains were stable and detected in at least three out of four timepoints, but one belonging to order Clostridiales was not detected consistently. These findings were similar at the 5-year time interval, where 43/48 original strains were detected while four strains belonging to order Clostridiales and one strain belonging to Lactobacillales were not present (**Figure 2D**). Since most of these were also not consistently detected in all the weekly timepoints, it is likely that they might be present at lower abundance and below the threshold for detection. Overall, these results confirm prior observations of long-term stability of the human gut microbiota using our sensitive and quantitative strain detection approach.

To determine if a similar long-term engraftment occurred in the non-relapsing post-FMT gut microbiota, we tracked 10 recipients for up to five years after FMT and found consistently high engraftment of donor strains at all time points (combined in **Figure 2E**, individual trajectories for donor-recipient pairs in **Supplementary Figure 2C**). In these individuals, we report an average engraftment of 83% (sd = 9%) at 36 hours, which stabilizes at 71% (sd = 16%) at 8 weeks and remains consistently high at 71% (sd = 9%) even 5 years later. In **Figure 2D** where a single donor was used for FMT in multiple individuals, we found similar engraftment of 79% (38 of 48 strains engrafted) at 8 weeks in a recipient which remained stable at 75% (36 out of 48 donor strains) 5 years later.

We found 50 out of 51 strains belonging to order Bacteriodales which engrafted at 8-weeks to remain stably engrafted for 6-months or more (**Supplementary Figure 2D**). However, fewer strains belonging to order Bifidobacteriales, which engrafted at 8-weeks remained stably engrafted at 6-months or longer timescale (only 5 out of 11, p-val < 10^−5^ fisher-exact test). Overall these results demonstrate that gut microbiota manipulation by FMT can lead to a near permanent engraftment of a new stable set of bacterial strains in patients with rCDI.

### FMT results in loss of original resident strains

Several studies have shown that resident microbiota strains create ecological niches^57,58^, which in turn can influence the engraftment of other microbes post-FMT. Thus, it is critical to identify the bacterial strains present pre-FMT and resolve their persistence dynamics after transplantation. Therefore, we isolated and sequenced the pre-FMT resident strains in 7 recipients, and tracked them for up to 5 years in each recipient’s metagenome. Similar to the PED metric, we defined Proportion Persistence of Recipient Strains (PPR) as the ratio between the strains of the recipient observed post-FMT to total recipient strains cultured before. Unlike the rapid high engraftment of donor strains, we found a more graduated decline in the PPR (combined in **Figure 3A**, individual trajectories for donor-recipient pairs in **Supplementary Figure 3A**) with the overall persistence decreasing to 49% (sd = 28%) at 1 week and 21% (sd = 10%) at 8 weeks (P val < 0.02 from one side Wilcox test). The recipient strains belonging to order Bifidobacteriales always persisted (7 of 7) in the recipients for 8 weeks post-FMT (**Supplementary Figure 3B, 3C**). However, recipient strains from order Lactobacillales and Enterobacterales were largely eliminated by the FMT. As in previous studies^59^, we observe an instability of the recipient’s gut microbiota where most resident strains of the recipient are lost after therapy, while the high sensitivity and resolution of our approach demonstrate that a focused subset of the original strains durably remain.

**Figure 3.**
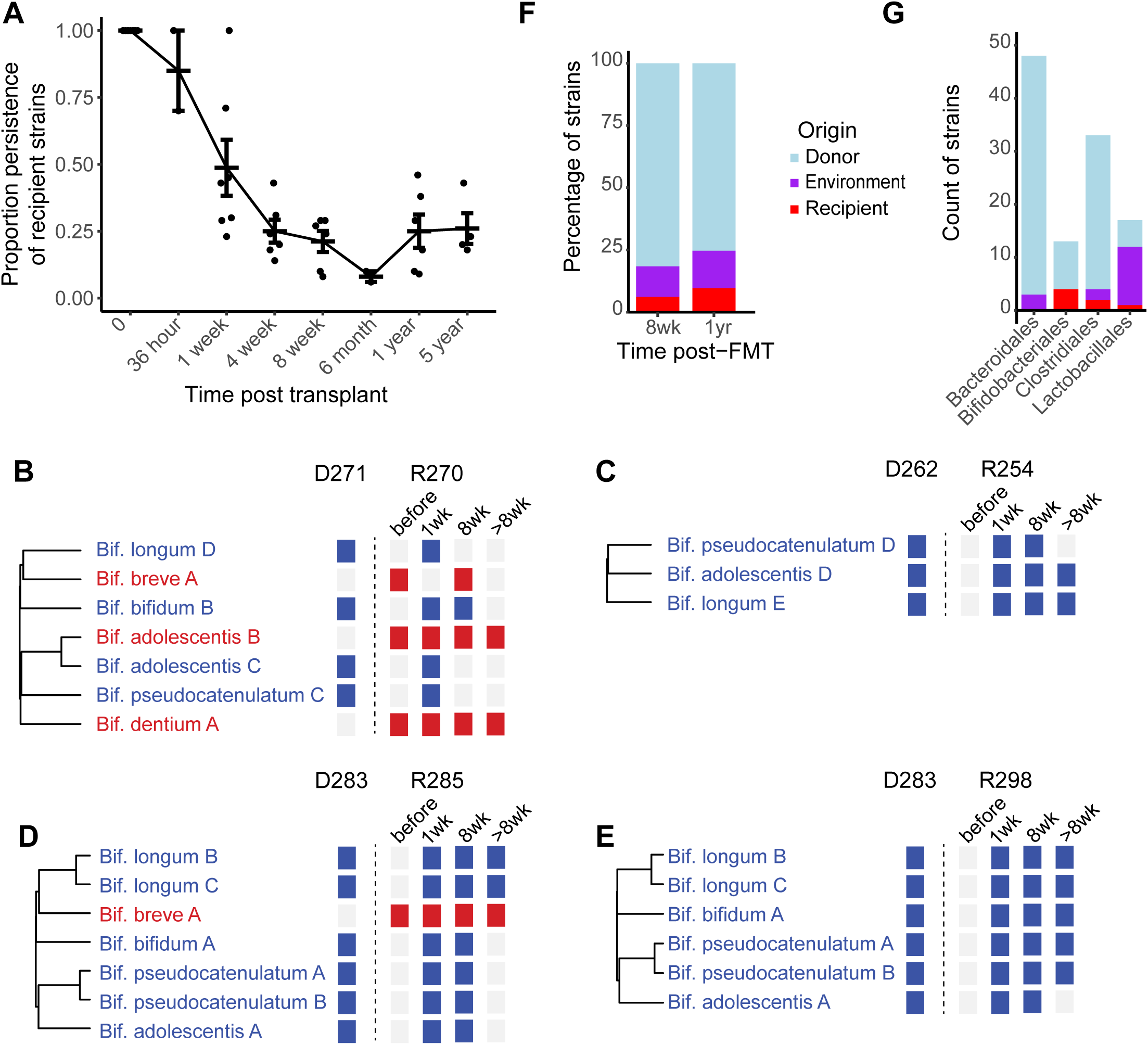
Persistence of recipient’s microbiota and engraftment of novel strains from the environment post-FMT. (**A**) Strains isolated from a recipient prior to FMT are rapidly lost with a small proportion persisting at longer timescales. (**B, C, D, E**) Strains belonging to order Bifidobacteriales in a recipient (red) prior to FMT, are associated with more limited engraftment of similar taxa from the donor (blue). Each column indicates a metagenomic sample. If a strain is detected in a metagenomics sample, it is colored based on origin of the strain. (**F**) Proportion of donor, recipient, and environment strains detected in patients post-FMT at both short (8-weeks) and longer time scales (1-year). Environmental strains are non-donor and non-recipient (prior to FMT) in origin, which are both cultured and metagenomically detected at that timepoint. (**G**) Count of strains detected in patients post-FMT at 8-weeks, subclassified by major phylogenetic taxa (at order level) and colored based on their origin.

### Recipient strains predict the engraftment of similar taxa from the donor

Earlier we observed lower engraftment of donor strains from the order Bifidobacteriales in recipients at 8 weeks (**Supplementary Figure 2B**) which further reduced at longer time scales (**Supplementary Figure 2D**). In contrast, the persistence of recipient’s original strains from the order Bifidobacteriales was consistently high after FMT (**Supplementary Figure 3C**). We therefore investigated if the recipient strains had a potential influence on the engraftment of similar taxa from the donor microbiota. In **Figures 3B-3E**, we present 4 case studies, where the presence of strains from recipient belonging to order Bifidobacteriales anticipate the relatively poorer engraftment of strains from this taxon from the donor. In Study 1 (**Figure 3B**), and the only FMT intervention where the recipient had multiple isolated strains (n = 3) belonging to order Bifidobacteriales, we observe that only 1 out of 4 Bifidobacteriales strains from the donor engraft at 8 weeks, while none engraft at longer time scales of 6 months or more. In contrast, all 3 strains from the recipient persist at 8 weeks and 2 out of 3 persist at longer time scale. In Study 2 (**Figure 3C**), the recipient had no strain belonging this order and all 3 strains from the donor are present at 8 weeks and 2 out of 3 are present at longer time scale.

In Studies 3 and 4, we focused on a single donor D283 which provided its sample for multiple FMT recipients (**Figure 2D**) including patients R285 and R298 which did not relapse and had samples collected at distant time points. Patient 1285 (Study 3, **Figure 3D**) had a single strain from order Bifidobacteriales which remained engrafted at both short and longer time scale. However, all 6 strains from the donor were present at 8 weeks, but reduced to 2 out of 6 at a longer time point. Patient 1298 (Study 4, **Figure 3E**) had no strains from this order and in contrast all 6 strains from the donor were present at 8 weeks, and only reduced to 5 at longer time scale. These results suggest that particular taxonomic groups^60^ in the recipient prior to FMT can limit the engraftment of specific taxonomic groups from the donor.

### Engraftment of non-donor strains after FMT

FMT results in majority engraftment of donor strains and majority loss of original recipient strains post-intervention. Given the unique physiology and environment of each individual, it is unlikely that this combination of events leads to complete niche occupancy of the host. Therefore, post-transplant there is likely at least a brief window where the recipient is most receptive to further engraftment of gut microbes from family members, other individuals and environmental sources of gut microbes. To test this hypothesis, we isolated and tracked strains from 4 patients post-FMT (at 8 weeks or 1 year). Together, we found 28 strains (**Supplementary Figure 3D**) that were non-donor and non-recipient in origin that were metagenomically detected and cultured in recipients post-FMT. On average in a patient post-FMT, 7.7% strains persisted from the recipient pre-FMT, 78.8% strains engrafted from the donor, and 13.5% strains were non-donor or non-recipient in origin (**Figure 3F**). This proportion was consistent across both short (8 weeks) and longer time scale (1 year). Although their origin and mode of transfer remains unknown, these environmental strains belong to phylogenetic taxa detected in both healthy donors and recipients prior to FMT (**Figure 3G**). We also found similar patterns in colonization of these environmental strains, where strains belonging to order Bacteriodales and Bifidobacteriales were always stable and detected at different timepoints, while those belonging to order Lactobacillales (only 7 out of 16 remained stable) often appeared at a timepoint but did not stably colonize (**Supplementary Figure 3D**). These results suggest that approximately 13.5% of the recipient niche space is unoccupied and stably colonized by other sources and suggest that LBPs with likely more limited niche occupancy will require a larger acquisition of environmental microbes for the host to become fully colonized.

### Donor engraftment independently explains rCDI FMT clinical outcomes

FMT which has a high success rate in treating patients with rCDI has no established metric to predict relapse or other clinical events. We have previously shown that patients with rCDI who have an early relapse have lower engraftment of donor strains at 8-weeks (**Figure 2B**), but it is not clear if engraftment metrics can elucidate later relapse or outcome of repeat-FMT in patients. To understand if our quantification of donor engraftment can independently explain the FMT outcome at a finer timescale, we tracked the proportional engraftment of donor strains in 5 patients that reported a relapse after FMT. In the three patients with stool samples collected near the date of relapse, we found engraftment to be low before or just after the relapse was noticed clinically (**Figure 4**). Patient R311 (top-left) had a low engraftment (8%) of donor microbiota strains at day 7 post-FMT and clinical relapse was detected on day 29. For patient R095 (top-right), the engraftment reduced moderately (54% to 46%) on day 32 post-FMT but reduced substantially (17%) on day 70, suggesting a clinical relapse closer to that date. Later by independently accessing the clinical records for this patient, we noticed that clinical relapse was diagnosed on day 65. Patient R216 (bottom-left) had high engraftment (75%) at day 44 post-FMT but low (13%) in the next time point collected on day 391. The clinical records revealed no-relapse on day 44, when our approach detected a high engraftment, but a relapse on day 90 which our approach detected at the next available timepoint on day 391 (engraftment of 13% down from 75%).

**Figure 4.**
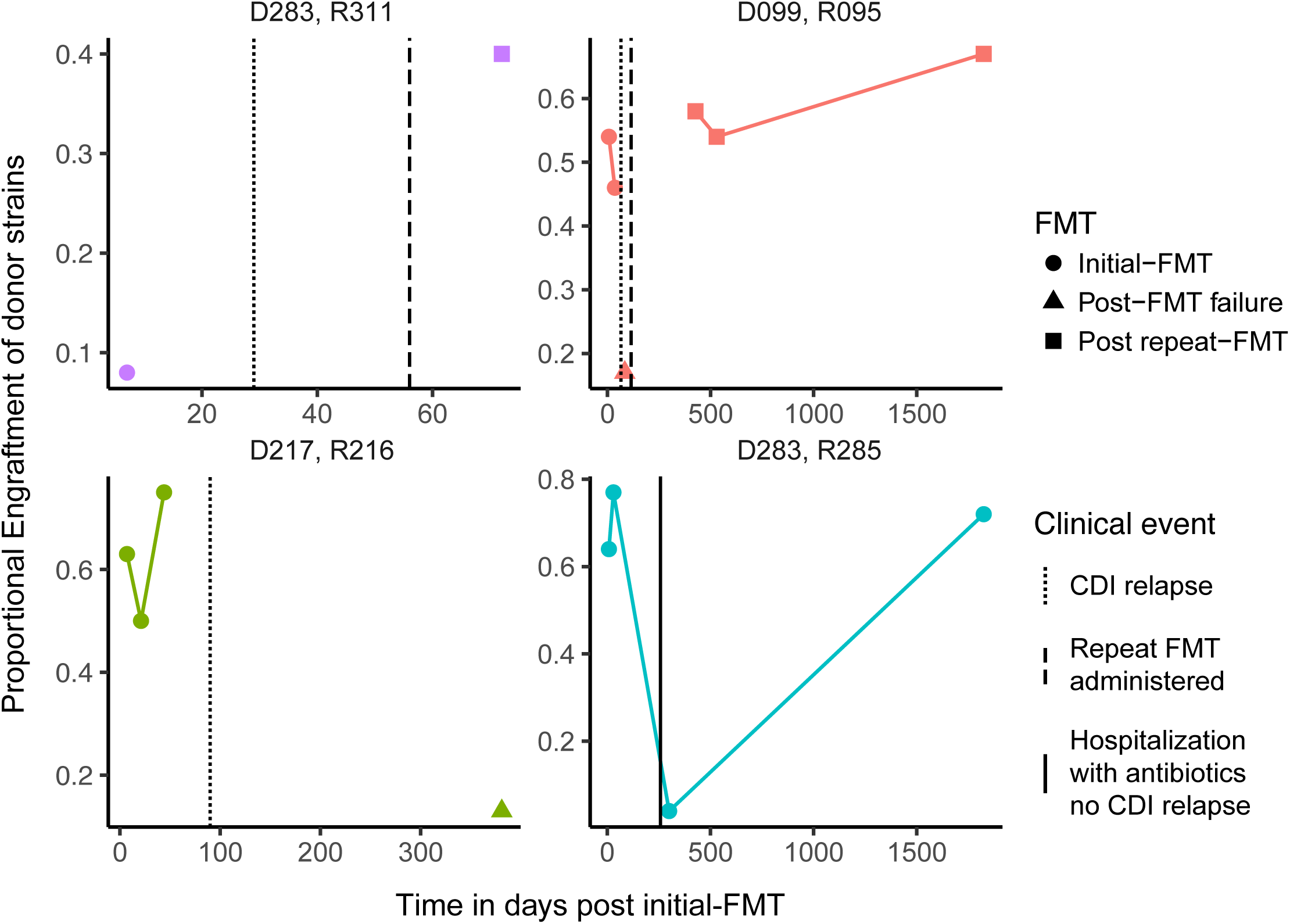
Donor strain engraftment in a recipient can predict FMT success or relapse (even after repeat FMT). Each individual trajectory represents the engraftment of donor strains in that recipient. Clinical events (noted independently of metagenomics) are indicated by differently shaped vertical lines. Note that R285 did not relapse but rather had a temporary loss in detectability of the donor strains during antibiotic treatment for severe diarrhea. The engrafted community fully recovered without FMT following the episode. R095 and R311 received a repeat FMT from the same donor, while R216 received an FMT from a different donor outside of this study so we were unable to quantify the long-term engraftment for this recipient.

Patients R095 and R311 received a repeat-FMT (from the same earlier donor) after the initial failure, and we detected a significant increase in strain engraftment in these recipients after the second FMT. They also remained relapse-free for the duration of the sampling period for up to 5 years.

For patient R285 (bottom-right), which had a successful FMT and reported high engraftment of 77% at day 30 post-FMT, the engraftment was reduced to 4% on day 300 but increased again 5 years later to 72%. The patient was symptom free at both 2 months and 5 years, in sync with expectations due to higher engraftment at those timepoints. Later by independently accessing the clinical records, we noticed that this patient was hospitalized with severe diarrhea and antibiotics on day 258 post-FMT, which our algorithm detected independently using metagenomics data collected on day 300. Importantly, this individual was not given a repeat FMT, suggesting the lower engraftment at day 30 post-FMT resulted in the large majority of engrafted strains being reduced below the detection limit of our algorithm but impressively not being eliminated from the gut.

Together, these results suggest that engraftment of donor strains at any time point is an accurate and robust metric for independently explaining the clinical outcome of FMT, both for initial and after a repeat FMT. However, the reasons for lower engraftment resulting in unsuccessful FMT in patients still remain unclear.

## Discussion

We developed the *Strainer* algorithm to track sequenced bacterial strains in low depth metagenomic sequencing. In combination with high throughput strain culturing and metagenomic sequencing of 14 donor and recipient pairs over multiple timepoints, we determined the majority of FMT donor strains (>70%) and a minority of recipient strains (<25%) are retained for at least five years after the transplant. The final recipient was composed of strains derived approximately 80%, 10%, and 10% from the donor, recipient, and environment respectively. We identified the proportional engraftment of donor strains is a predictive measure of FMT success and rCDI relapse with all non-relapsing subjects having donor engraftment >35% at all time points and all relapsing subjects having PED <20% at samples near the time of relapse. While successful FMT is associated with high engraftment of donor strains, this engraftment is driven by strains from a subset of taxonomic groups that engraft very well, in line with previous studies^61^ that suggest the role of specific group of strains in determining the long-term outcome of FMT intervention in patients.

We acknowledge that our approach is dependent on the sequencing of bacterial strains, which it detects and tracks from the metagenomes. Approaches for high throughput strain culturing and isolation are improving but are still limited by factors of cost, time and difficulty in culturing of microbes from some species. Our approach is most suited for situations when one donor or LBP is transplanted into many recipients, which will likely be the dominant use-case if LBPs equivalent to FMT are developed for some clinical indications. In addition, advances in the scale of long-read sequencing will potentially lead to greatly improved metagenomic assemblies from single fecal samples^62^, which could similarly be used by our approach.

The quantitative nature of our framework can be used to inform gut microbiota manipulation study design beyond clinical efficacy to optimize the conditions for the durable structural modification of the host microbiome. As an example, if oral FMT capsules result in similar engraftment and outcomes as colonoscopic infusion of donor material^20,21^, future study designs may prefer the less invasive oral approach. Tracking differences in engraftment between approaches that do not differ in clinical efficacy could help narrow our knowledge of what strains are necessary for an LBP alternative to FMT. Some FMT trials^22,24^ used multiple transplants over many weeks, and our framework and algorithm could accurately evaluate the engraftment gains to determine the cost/benefit of repeated doses.

Recently, the FDA^63–65^ has issued multiple safety alerts regarding the use of FMT and risk for serious adverse events due to transmission^66^ of multi-drug resistant organisms (MDRO) in the fecal material used for FMT. An important result from this study is the identification of a select mixture of live bacterial strains that stably engraft for 5 years in patients post successful and symptom free FMT. These strains are an ideal starting point for a synthetic FMT alternative for rCDI of validated engrafting strains free of MDROs. Moreover, our approach is well-placed to track the engraftment of such a cocktail in shallow metagenomic sequencing from hundreds to thousands of patients to monitor the long-term success of FMT.

In summary, our sensitive algorithm and long-term clinical sampling of FMT recipients have revealed that the majority of FMT donor bacteria engraft for at least five years in non-relapsing individuals demonstrating that gut microbiota manipulation represents a uniquely durable therapeutic paradigm.

## Supplementary Figures

**Supplementary Figure 1.**
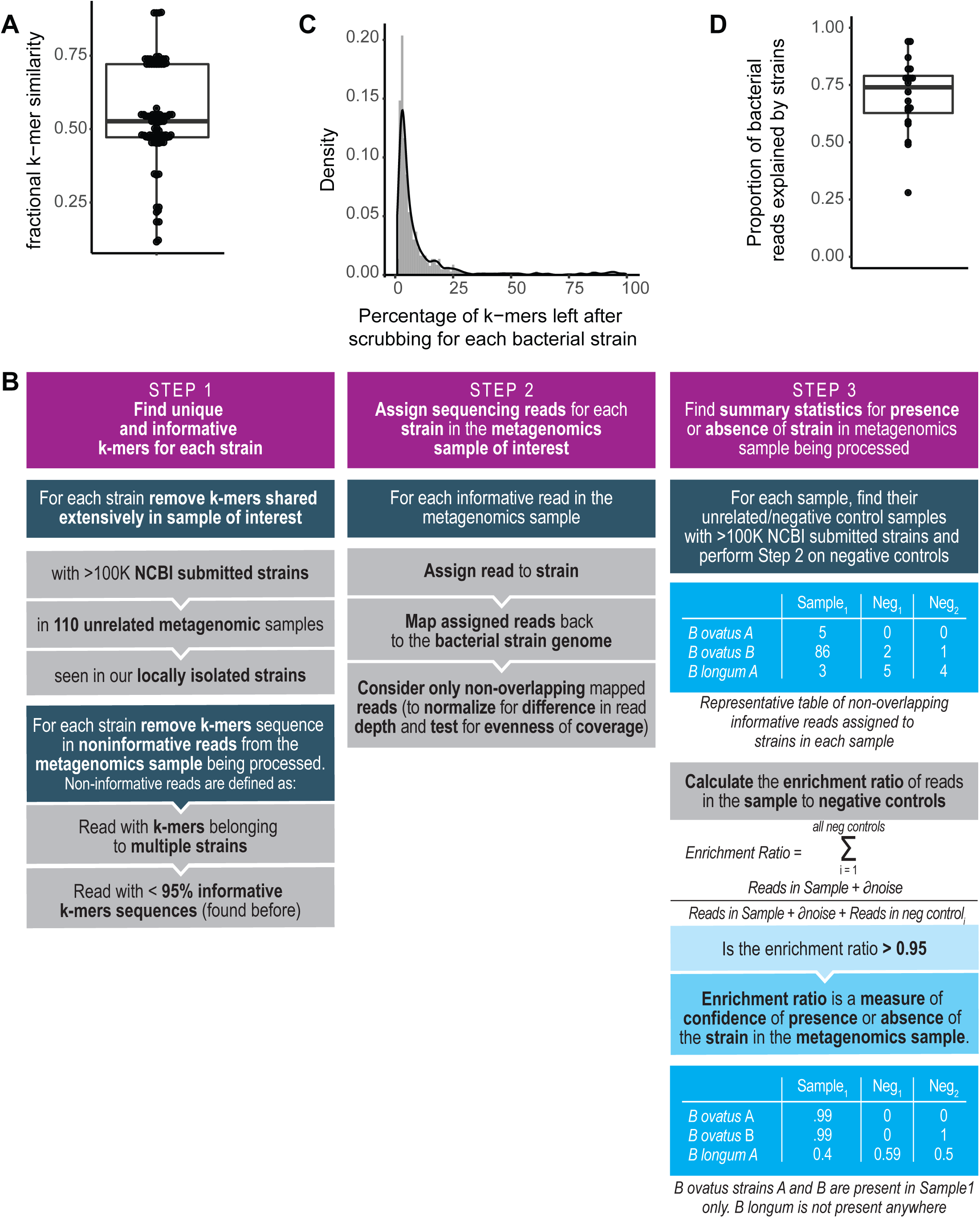
The *Strainer* algorithm. (**A**) Percentage similarity between 96 different isolates of species *Bacteriodes ovatus* and the reference strain AAXF00000000.2. Similarity is found by comparing sequence k-mers of length 31 between genomes. (**B**) Overview of our algorithm *Strainer*. The algorithm has 3 modules, where Module-1 involves finding the unique and likely informative sequence k-mers for each strain by removing those shared extensively with unrelated sequenced strains in NCBI, unrelated metagenomics samples, and those cultured and sequenced in this study. Next, we decompose each sequencing read in the metagenomics sample of interest into its k-mers, and find reads which have k-mers belonging to multiple strains, or have <95% of informative k-mers for a single strain. We further remove these non-informative k-mers from our previous set. In Module-2 we assign sequencing reads from the metagenomics sample of interest, with a majority of informative k-mers (>95%) to each strain. Next, we map these reads to the genome of the corresponding strain, and consider the non-overlapping ones only. This step normalizes for sequencing depth across samples and checks for evenness of read distribution across the bacterial genome. Finally, in Module-3 we compare the read enrichment in a sample to unrelated samples or negative controls and present summary statistics for presence or absence of a strain in a sample. (**C**) Density plot of percentage of k-mers remaining for each strain, after removing those shared extensively with other unrelated bacterial genomes and metagenomics samples. (**D**) Proportion of bacterial reads in the metagenomics sample that are explained by the genome sequences of the cultured strain library for that sample. Each point in the boxplot corresponds to a separate sample.

**Supplementary Figure 2.**
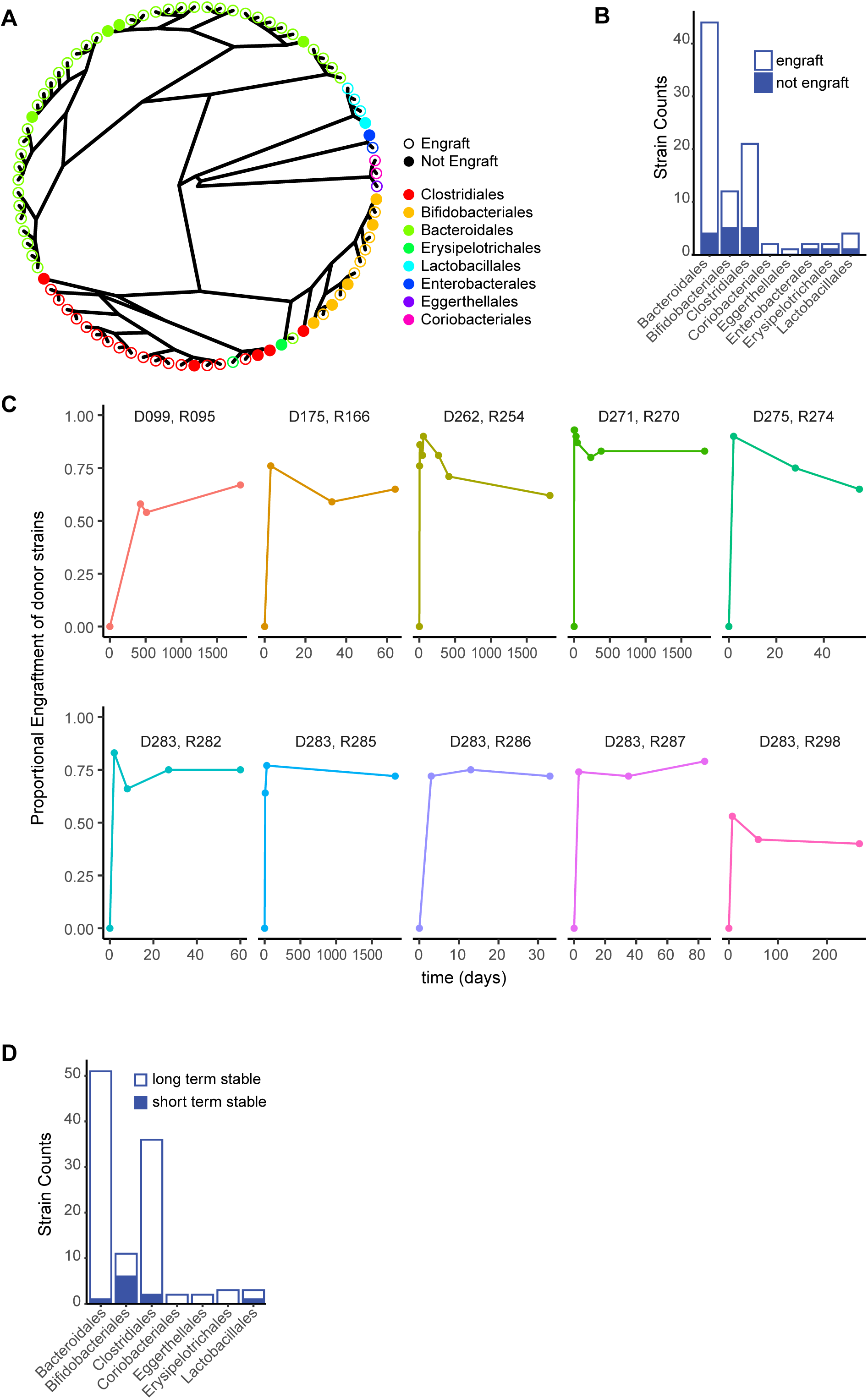
Donors bacterial strains engraftment in recipients post-FMT. (**A**) Radial phylogeny plot identifying bacterial strains that engraft for at least 8-weeks in patients post-FMT. Filled circle indicates the strain did not engraft. The colors represent different taxonomic orders. (**B**) Number of strains that transmit and engraft for at least 8-weeks in patients post-FMT (single FMT donor to recipient setting) grouped by taxonomic order. (**C**) Trajectory of proportional strain engraftment of donor strains in each recipient at all available timepoints (in days). The donor recipient pair ids are at the top of each plot. (**D**) The number of strains colonized at 8 weeks (short term) that engraft for at least 6-months or more (long-term) in patients post-FMT (both single FMT donor to single and multiple recipients setting) grouped by taxonomic order.

**Supplementary Figure 3.**
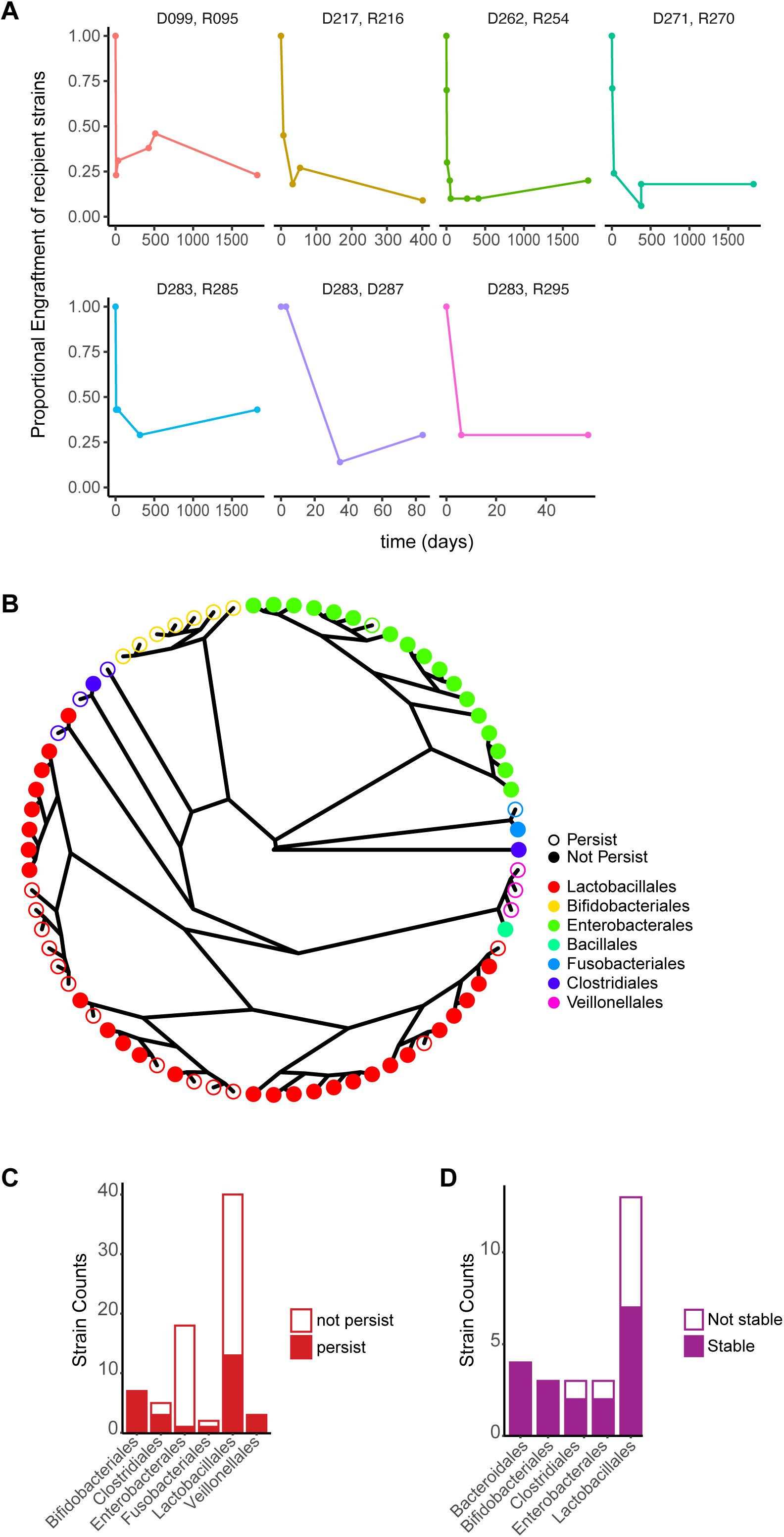
Recipient strains persistence and engraftment of novel strains from the environment. (**A**) Trajectory of proportional persistence of recipient’s strains post-FMT at all available timepoints (in days). The donor recipient pair ids are at the top of each plot. (**B**) Radial phylogeny plot identifying original recipient’s bacterial strains that persist for at least 8-weeks post-FMT. Filled circle indicates the strain did not persist. The colors represent different taxonomic orders. (**C**) The number of the recipient’s original strains that persist for at least 8-weeks post-FMT, grouped by taxonomic order. (**D**) The number of environment strains (i.e. non-donor and non-recipient in origin) that engraft in patients stably over multiple timepoints (>1 week) post-FMT, grouped by taxonomic order.

## Supplementary Tables

**Supplementary Table 1**. Bacterial strains, their phylogenetic classification, accession numbers and the percentage of k-mers remaining after removing previously seen k-mers.

**Supplementary Table 2**. *Bacteroides ovatus* strains and details on their sequential colonization in gnotobiotic mice. This truth table of the presence (1) or absence (0) of each strain in a sample, which is presented in Figure 1A.

**Supplementary Table 3**. Truth table for presence (1) or absence (0) of each strain in 10 healthy donors, which is presented in Figures 1B and C. Presence (1) of strain implies that the strain was isolated and cultured from that sample.

**Supplementary Table 4**. Truth table for presence (1) or absence (0) of each strain in 5 patients with rCDI, which is presented in Figure 1C. Presence (1) of strain implies that the strain was isolated and cultured from that sample.

**Supplementary Table 5**. Truth table for presence (1) or absence (0) of each strain in 4 patients with IBD, which is presented in Figure 1C. Presence (1) of strain implies that the strain was isolated and cultured from that sample.

**Supplementary Table 6**. All metagenomics samples and their accession numbers.

**Supplementary Table 7**. Details for each FMT, samples collected at multiple timepoints, details of strains isolated at that timepoint (if any) and other clinical information (relapse, non-relapse or hospitalization status).

## Methods and Materials

### Germ-free mice and colonization with cultured bacteria

The mice experiments were performed as a part of other published studies for understanding the strain-level differences and their role in fecal-IgA levels^7,67,68^, as well as the impact of the IBD and non-IBD microbiome on baseline immune tone and colitis. These data were used to evaluate the performance of our *Strainer* algorithm in defined communities of bacterial strains in germ-free mice.

### Human subjects

All individuals of age 18 and over were recruited in the study using a protocol approved by the Mount Sinai Institutional Review Board (HS# 11-01669). The donors and patients who received FMT for rCDI or rCDI and IBD were described in a previous study analyzed with 16S rRNA amplicon sequencing^69^.

### Fecal sample collection, DNA extraction and shotgun metagenomic sequencing

We followed the protocol previously described in our published^12,46,48^ studies. Briefly, samples were aliquoted on dry ice or liquid nitrogen and stored at -80C. DNA was then extracted by bead beating in phenol chloroform. Illumina sequencing libraries were generated from sonicated DNA, ligation products purified and finally enrichment PCR performed. Samples were pooled in equal proportions and size-selected before sequencing with an Illumina HiSeq (paired-end 150 bp). Sequence data files (fastq) for all metagenomic sequencing samples are stored in the public Sequence Read Archive (SRA) under project number PRJNA637878.

### High-throughput anaerobic bacterial culture

We utilize a well-established, robotized platform that enables isolation and culturing of a high proportion of the bacteria found in the human gut^12,39,48,68^. Briefly, the steps involve first plating clarified stool samples on solid media, followed by growth under a range of environmental conditions designed to cultivate anaerobic, microaerophilic, aerobic and spore-forming bacteria. Next, 384 colonies are picked for each donor sample and regrown in liquid media in multiwell plates. Each isolate is then identified by a combination of MALDI-TOF mass spectrometry, 16S rDNA and whole genome sequencing. Using this knowledge, the original 384 isolates are de-replicated and unique strains for each donor are archived in multiwell plates, which allows for automated selection of specific strains and sub-communities. The sequencing reads from each cultured isolated were quality filtered with Trimmomatic^70^ and assembled using Spades^71^.

### Strain as bacterial isolates with <96% similarity

Similar to our previous analyses^12,39,48^, bacterial isolates with <96% whole genome similarity were defined as unique strains, otherwise they were considered as multiple isolates for the representative strain. Pairwise strain similarity was found with k-mer counting software kmc version 3.0^72^

### Strainer framework for detection of bacterial strains from metagenomic samples

The algorithmic framework *Strainer* has 3 separate parts (**Supplementary Figure 1B**), where the first part involves finding the unique and likely informative sequence k-mers (k = 31) for each strain by removing those shared extensively with unrelated sequenced strains in NCBI, unrelated metagenomics samples, and those cultured and sequenced in this study. Any k-mer shared two times or more with these unrelated samples is marked for removal. However, some bacterial species share genomic sequences extensively and this stringent criterion is relaxed (i.e. sharing cutoff is iteratively increased) until we have at least 4% of genomic k-mers left for each bacterial strain. Next, we decompose each sequencing read in the metagenomics sample of interest into its k-mers, and find reads which have k-mers belonging to multiple strains, or have <95% of informative k-mers for a single strain. We further remove these non-informative k-mers from our previous set. In the second part we assign sequencing reads from the metagenomics sample of interest, with a majority of informative k-mers (>95% found at the end of part-1) to each strain. Next, we map these reads to the genome of the corresponding strain (bowtie2^73^ with very-sensitive no-mixed no-discordant options), and consider the non-overlapping ones only. This step normalizes for sequencing depth across samples and checks for evenness of read distribution across the bacterial genome. Finally, in the last part we compare the read enrichment in a sample to unrelated samples or negative controls and present summary statistics for presence or absence of a strain in a sample.

### Statistical analysis and plotting

Precision Recall curve were plotted with R package PRROC version 1.3.1. Analysis was performed in Python version 2.7.16 and R studio version 1.1.453. Radial graph for phylogeny were plotted using R packages ape version 5.4 and phytools version 0.7. Our statistical framework, associated databases and code is publicly available. The authors would enthusiastically respond to all reasonable requests for customization of *Strainer* code and statistical framework.

## Abbreviations used

AUC: Area Under the Curve
Blu: Blautia
Rum: Ruminococcus
Bif: Bifidobacterium
Bac: Bacteriodes
Esc: Escherichia
Ali: Alistipes
Par: Parabacteriodes
Bar: Barnesiella
Clo: Clostridium
Eub: Eubacterium
Pho: Phocea
Cop: Coprococcus
Egg: Eggerthella
Ery: Erysipelatoclostridium
Dor: Dorea
Ros: Roseburia
Odo: Odoribacter
Par: Parabacteriodes
Str: Streptococcus

## Acknowledgements

This work was supported in part by the staff and resources of the Microbiome Translational Center and the Scientific Computing Division in Icahn School of Medicine at Mount Sinai. We thank C. Fermin, E. Vazquez, and G.N. Escano for gnotobiotic husbandry support and S. Simmons for helpful suggestions. This work was supported by National Institutes of Health Grants (NCCIH AT008661, NIDDK DK112978, NIDDK DK124133, SUCCESS), and Crohn’s and Colitis Foundation RFA awards 650451 and 580924.

## Author Contributions

V.A. and J.J.F wrote the manuscript. I.M, Z.L, C.Y, G.J.B and A.CL collected samples and also performed experiments. J.M and A.G collected the clinical samples. V.A, J.J.F, I.M, C.Y, G.J.B, A.CL, A.G, D.G, J.C.C and J-F.C were involved in data analysis and interpretation. All authors read, provided critical feedback and approved the final manuscript.

